# Asymmetric Contraction of Adherens Junctions arises through RhoA and E-cadherin feedback

**DOI:** 10.1101/2021.02.26.433093

**Authors:** Kate E. Cavanaugh, Michael Staddon, Theresa A. Chmiel, Robert Harmon, Srikanth Budnar, Alpha S. Yap, Shiladitya Banerjee, Margaret L. Gardel

**Affiliations:** Committee on Development, Regeneration, and Stem Cell Biology, University of Chicago, Chicago, Illinois 60637, USA; James Franck Institute, Department of Physics, Pritzker School of Molecular Engineering, University of Chicago, Chicago, Illinois 60637, USA; Institute for Biophysical Dynamics, Pritzker School of Molecular Engineering, University of Chicago, Chicago Illinois 60637, USA; Center for Systems Biology Dresden, 01307 Dresden, Germany; Division of Cell and Developmental Biology, Institute for Molecular Bioscience, The University of Queensland, St. Lucia, Brisbane, QLD 4072, Australia; CSL Ltd, Bio21 Institute, Melbourne, Victoria 3052, Australia; Department of Physics, Carnegie Mellon University, Pittsburgh, PA 15213, USA

## Abstract

Tissue morphogenesis often arises from the culmination of discrete changes in cell-cell junction behaviors, namely ratcheted junction contractions that lead to collective cellular rearrangements. Mechanochemical signaling in the form of RhoA underlies these ratcheted contractions, which occur asymmetrically as one highly motile vertex contracts toward a relatively less motile tricellular vertex. The underlying mechanisms driving asymmetric vertex movement remains unknown. Here, we use optogenetically controlled RhoA in model epithelia together with biophysical modeling to uncover the mechanism lending to asymmetric vertex motion. We find that both local and global RhoA activation leads to increases in junctional tension, thereby facilitating vertex motion. RhoA activation occurs in discrete regions along the junction and is skewed towards the less-motile vertex. At these less-motile vertices, E-cadherin acts as an opposing factor to limit vertex motion through increased frictional drag. Surprisingly, we uncover a feedback loop between RhoA and E-cadherin, as regional optogenetic activation of specified junctional zones pools E-cadherin to the location of RhoA activation. Incorporating this circuit into a mathematical model, we find that a positive feedback between RhoA-mediated tension and E-cadherin-induced frictional drag on tricellular vertices recapitulates experimental data. As such, the location of RhoA determines which vertex is under high tension, pooling E-cadherin and increasing the frictional load at the tricellular vertex to limit its motion. This feedback drives a tension-dependent intercellular “clutch” at tricellular vertices which stabilizes vertex motion upon tensional load.

## Introduction

Morphogenesis relies on the tight spatiotemporal control of cell-cell junction lengths (Lecuit et al., 2011). Contractile forces, acting at adherens junctions, alter junction lengths as a cyclic ratchet (Fernandez-Gonzalez and Zallen, 2011; Jewett et al., 2017; Mason et al., 2013; Rauzi et al., 2010; Solon et al., 2009). Preceding these ratcheted contractions are pulses of active RhoA (Kerridge et al., 2016; Munjal et al., 2015; Rauzi et al., 2010), the strength and temporal pattern of which control junction tension to confer junction length (Cavanaugh et al., 2020; Staddon et al., 2019). Through effector activation, contractile actomyosin arrays assemble rapidly in response to intracellular biochemical signals and/or physical cues from neighboring cells (García-Mata and Burridge, 2007). As such, RhoA GTPase cycling is thought to give rise to spatiotemporal changes in junction length which, in turn, drive tissue morphogenesis(Mason et al., 2016). While the molecular components of these mechanochemical systems are well characterized, the mechanisms by which RhoA regulates junctional tension and adhesion to control cell shape remains largely unknown.

A recent study has revealed the asymmetric nature of junction contraction that occurs during germband extension (Vanderleest et al., 2018). Here, one tricellular vertex is highly mobile and contracts towards a more immobile, stationary vertex. The net result of this asymmetric vertex motion is coordinated asymmetry in junction deformation whose collective contractions facilitate global tissue rearrangements (Huebner et al., 2020a; Vanderleest et al., 2018). A possible mechanism underlying this innate vertex asymmetry describes heterogeneous force production along the junction proper. Non-uniform force production may cause very local actomyosin flows to specific regions of the junction for qualitatively different junctional responses. Bicellular edges, for example, act as independent contractile units apart from tricellular vertices (Choi et al., 2016; Vanderleest et al., 2018). Medioapical actomyosin flows to the bicellular interfaces can also generate contractile forces sufficient to deform junctions (Munjal et al., 2015; Rauzi et al., 2010). Flows to the tricellular vertices may restrict these contractions, thus stabilizing the junctional ratchet (Vanderleest et al., 2018). The coordination between these spatially distinct actomyosin flows may yield asymmetric junction shortening (Vanderleest et al., 2018). Thus, the molecular, cellular, and mechanical origins of this asymmetry remain unclear.

Cells sense and respond to mechanical cues through force-sensitive feedbacks within the cytoskeleton. Apical E-cadherin-based adhesions mediate intercellular cell-cell adhesion. However, E-cadherin should be envisaged not as a static participant of cellular adherence but rather as a dynamic sensor of force that dictates cellular behavior. For example, force stimulates the RhoA pathway and myosin light chain phosphorylation, resulting in an overall increase in actin polymerization at adherens junctions (Acharya et al., 2017). Additionally, force-sensitive processes within adherens junctions allows adhesive components to strengthen under force (Manibog et al., 2014). Here, cadherin catch bonds are strengthened when adhesion complexes experience tensile force (Buckley et al., 2014). Together these mechanisms cause clustering of E-cadherin molecules and actin to trigger adhesion complex growth (Hong et al., 2013). In this way, these proteins subsequently generate a reinforcement response to anchor junctions against applied force (Pannekoek et al., 2019). However, it is still unclear if and how cells’ force-sensitive coupling of actomyosin and adhesion complexes modulate junction length to coordinate morphogenetic movements within tissues.

Here, we investigated the origins of asymmetric junction contraction by using optogenetic and pharmacologic modulation of RhoA activity. We found that uniform RhoA activation along the junction drove asymmetric contraction. We then used computational modeling to offer predictions on the mechanistic origin of this asymmetric contraction. Our experimental data indicated that differential regulation of vertex tension, as predicted by canonical models of epithelial tissues, was insufficient to account for such asymmetry. We then explored how asymmetric RhoA-mediated contraction and asymmetric vertex friction accounted for experimental results. Here, we found that solely an asymmetry in RhoA localization generated qualitatively different junctional regimes of contraction and subsequent vertex asymmetry. Instead, we determined that force-dependent adhesion at tricellular vertices act to locally reinforce the vertex to restrict its movement. Feedback between RhoA-mediated tension and E-cadherin-induced friction was sufficient to recapitulate experimental data. By modulating E-cadherin friction with pharmacological perturbations, we induced symmetry back into the system or abolished junction contraction entirely. Our modeling and experimental data therefore point to a unified model of asymmetry induced by both friction and local contraction that is mediated by a RhoA-dependent mechanosensitive rigidity transition of tricellular vertices.

## Results

### RhoA stimulates Asymmetric Junction Contraction

To examine how RhoA controls junction contractions, we formed a model tissue by plating a colorectal adenocarcinoma (Caco-2) cell line endogenously CRISPR tagged for E-cadherin on polymerized collagen gels at full confluency to facilitate the monitoring of junctional movements (Figure 1A) (Liang et al., 2017). We then measured junction length by finding the interfacial distance from one tricellular vertex to the other tricellular vertex. In control conditions, there were negligible changes in junction length over the course of a two-hour period (Figure 1B, Supplemental Figure 1A, Supplemental Movie 1). Here, the junction length was maintained and only fluctuated about 1% over the two-hour period (Figure 1B).

**Figure 1:**
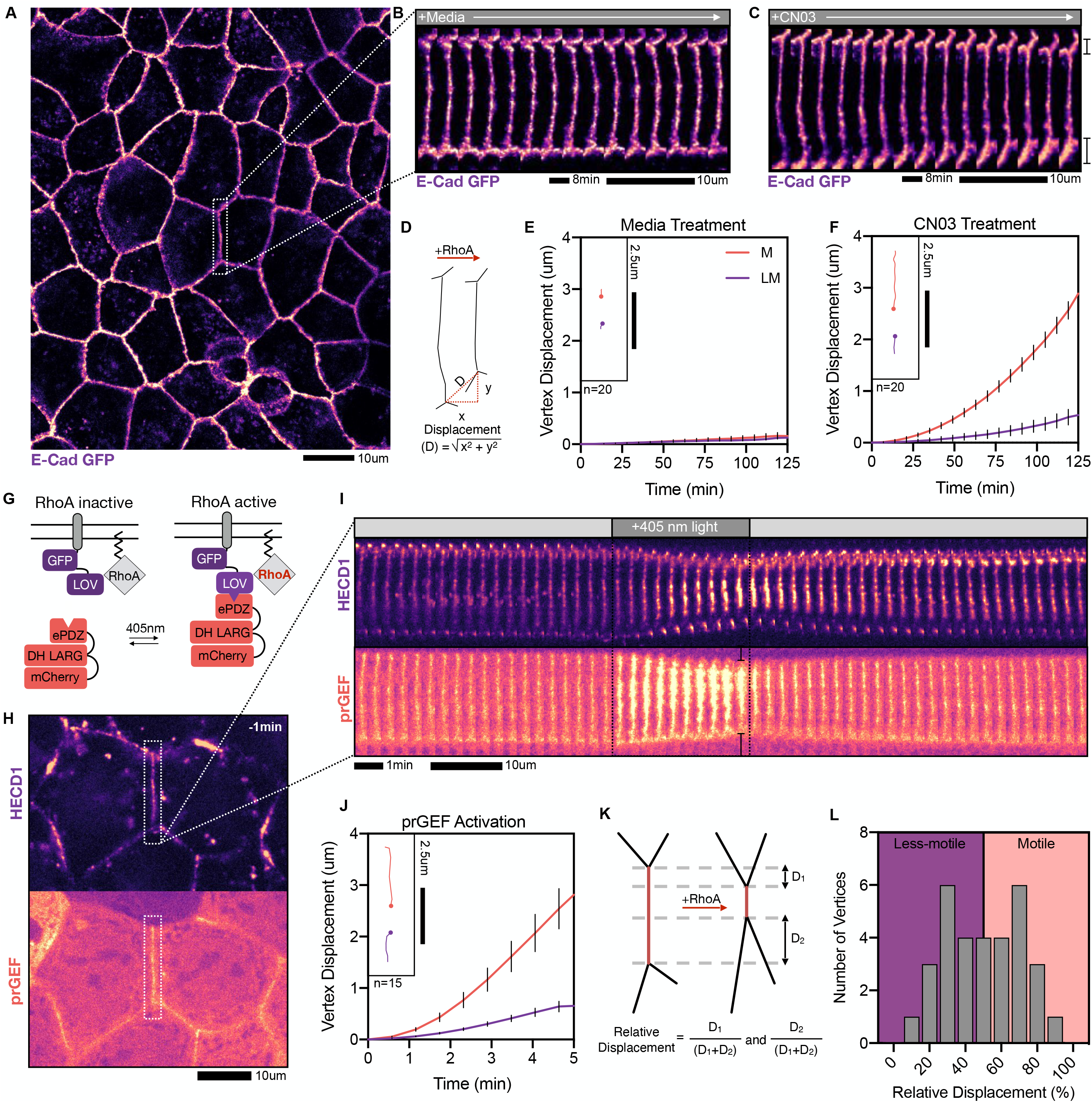
RhoA-mediated tension drives vertex motion. A. Representative image of a homeostatic epithelial tissue endogenously expressing E-Cadherin GFP. B. Zoomed in images of WT junction over the course of two hours showing no junction length changes with the addition of media. C. Representative images of timelapse video over the course of two hours showing asymmetric junction shortening with the addition of the CN03 compound. D. Schematic of junction shortening and displacement measurement analysis. E. Vertex displacement analysis for junctions in WT conditions showing little-to-no vertex motion. Inlay shows particle tracks for a representative vertex pair in WT conditions. F. Vertex displacement analysis for junctions in CN03 treatment showing asymmetry in vertex displacements. Inlay shows particle tracks for a representative vertex pair in CN03 treatment. G. Schematic of the TULIP optogenetic system to drive local RhoA activation. H. Zoomed out image of a targeted junction at −1min before optogenetic activation. Top image shows HECD1 junction labeling of E-cadherin and bottom image shows prGEF localization. I. Timelapse of the junction in H undergoing a 5-min optogenetic activation showing asymmetric junction contraction within the activation period and junction relaxation post-activation. J. Vertex displacement analysis for the junction within the 5-min optogenetic activation period. Displacement analysis shows similar vertex asymmetry upon increases in RhoA activity. Inlay shows particle tracks during the 5-min optogenetic activation period for a representative vertex pair. K. Schematic documenting the percent movement analysis. L. A histogram of the percent motions of all vertices undergoing optogenetic activation shows two peaks at 30% and 70%, further documenting vertex asymmetry.

We then treated cells with a cell permeable, pharmacological RhoA Activator, CN03, to globally and acutely increase RhoA activity across the entire tissue. We began imaging upon the addition of CN03, at time (t)=0 min, and examined junction fluctuations and contractions resulting from RhoA increases until (t)=125 min. About 30% of the junctions contracted, resulting in their shortening to about 80% of the initial length (Figure 1C, Supplemental Figure 1A, Supplemental Movie 1). We manually tracked each vertex over time and measured its displacement (Figure 1D). In control conditions, we found that there was little to no vertex movement (Figure 1B, 1E). In contrast, in CN03 containing media one vertex moved significantly more than the other vertex (Figure 1C, 1F). This asymmetric contraction is reminiscent of observations in developmental systems (Huebner et al., 2020b; Vanderleest et al.).

To explore the impact of localized Rho activity, we turned to an optogenetic approach. The logic behind this experiment was to have isolated junctions acutely experience heightened and targeted RhoA activation. For spatial and temporal control over RhoA activity, we used a Caco-2 cell line expressing the TULIP optogenetic two-component system (Cavanaugh et al., 2020; Oakes et al., 2017; Staddon et al., 2019; Strickland et al., 2012; Wagner and Glotzer, 2016). TULIP’s two components include the 1) membrane-tethered photosensitive LOVpep anchor protein and the 2) prGEF complex that houses the photorecruitable PDZ domain attached to the catalytic DH domain of the RhoA-specific GEF, LARG. Blue light (405nm) activation causes a conformational change in the LOVpep domain to expose a docking site for the engineered PDZ domain within the prGEF complex. This blue light activation increases the binding affinity between the two components, thereby recruiting the prGEF to the membrane where it drives local RhoA activation (Figure 1G) (Cavanaugh et al., 2020; Oakes et al., 2017; Staddon et al., 2019; Wagner and Glotzer, 2016). This system has high temporal resolution, as prGEF recruitment and dissociation occurs on the order of 30-60 seconds. prGEF recruitment was tightly confined to the targeted cell-cell junction, consistent with previously published work (Figure 1I) (Cavanaugh et al., 2020). Overall, this system gave tight spatiotemporal control over RhoA activity for which to study how junctions contract upon increased RhoA activity.

In order to visualize the movement of adhesive sites, serving as fiducial marks along the junction, we labeled E-Cadherin using an antibody labeling technique targeting its extracellular domain. We bathed the cells for at least an hour in E-cadherin primary antibody, HECD1, and its corresponding fluorescently labeled secondary antibody. Upon washing out the antibody, we found that this labeling produced a punctate pattern of E-cadherin that delineated the cell-cell junctions and vertices (Figure 1H, Supplemental Figure 1B). HECD1 targets the EC2 domain region of the E-cadherin ectodomain, rather than the EC1 domain which mediates trans-binding. In this way, cellular cohesion and intercellular E-cadherin binding via EC1 domains was preserved. Indeed, we found that under the conditions of our experiments, HECD1 did not affect junction contraction or length remodeling (Supplemental Figures 1C-E)(Cavanaugh et al., 2020). Specifically, we found that over a five-minute light activation period, the targeted HECD1-labeled junctions rapidly contracted to ~70-80% of their original length and then returned to their original lengths after light termination, consistent with previous reports (Figure 1I, Supplemental Figure 1C, 1D, Supplemental Movie 2) (Cavanaugh et al., 2020). This contraction was surprisingly consistent across multiple junctions with different initial lengths and geometries. Such asymmetry was also surprisingly unchanged by alterations to cell-substrate adhesions (Supplemental Figure 2A-I).

The vertex displacements in response to junctional RhoA activation were also asymmetric (Figure 1I, 1J, Supplemental Movie 2). To quantify the asymmetry, we measured relative displacement of each vertex in a vertex pair, as defined by the distance moved of one vertex (e.g. D_1_) over the total distance moved by both vertices (D_1_+D_2_), to yield D_1_/(D_1_+D_2_) and D_2_/(D_1_+D_2_) (Figure 1K). We then plotted the probability density of the relative movement. This revealed an asymmetry in the histogram with peaks around 30% and 70%, further indicating an inherent asymmetry in the distribution of vertex motion (Figure 1L). This result was starkly contrasted against a symmetric contraction, where a single peak centered around 50% would be expected. Interestingly, this vertex asymmetry occurred despite uniform prGEF recruitment along the junction, discarding the possibility that heterogeneous regions of RhoGEF recruitment could trigger asymmetric junction contraction (Supplemental Figure 1F, 1G).

### Asymmetric contraction can be driven by heterogeneity in RhoA activation

Junctions could either contract uniformly along their length or contain individual sub-junctional segments that contract varying amounts. To explore these possibilities, we used the variable intensities of HECD1 labeling to examine local variations in deformations along the junction. A line-scan along the junction, taken over time, created a kymograph for which to analyze fiduciary flows before, during, and after light-stimulated junction contraction (Figure 2A).

**Figure 2:**
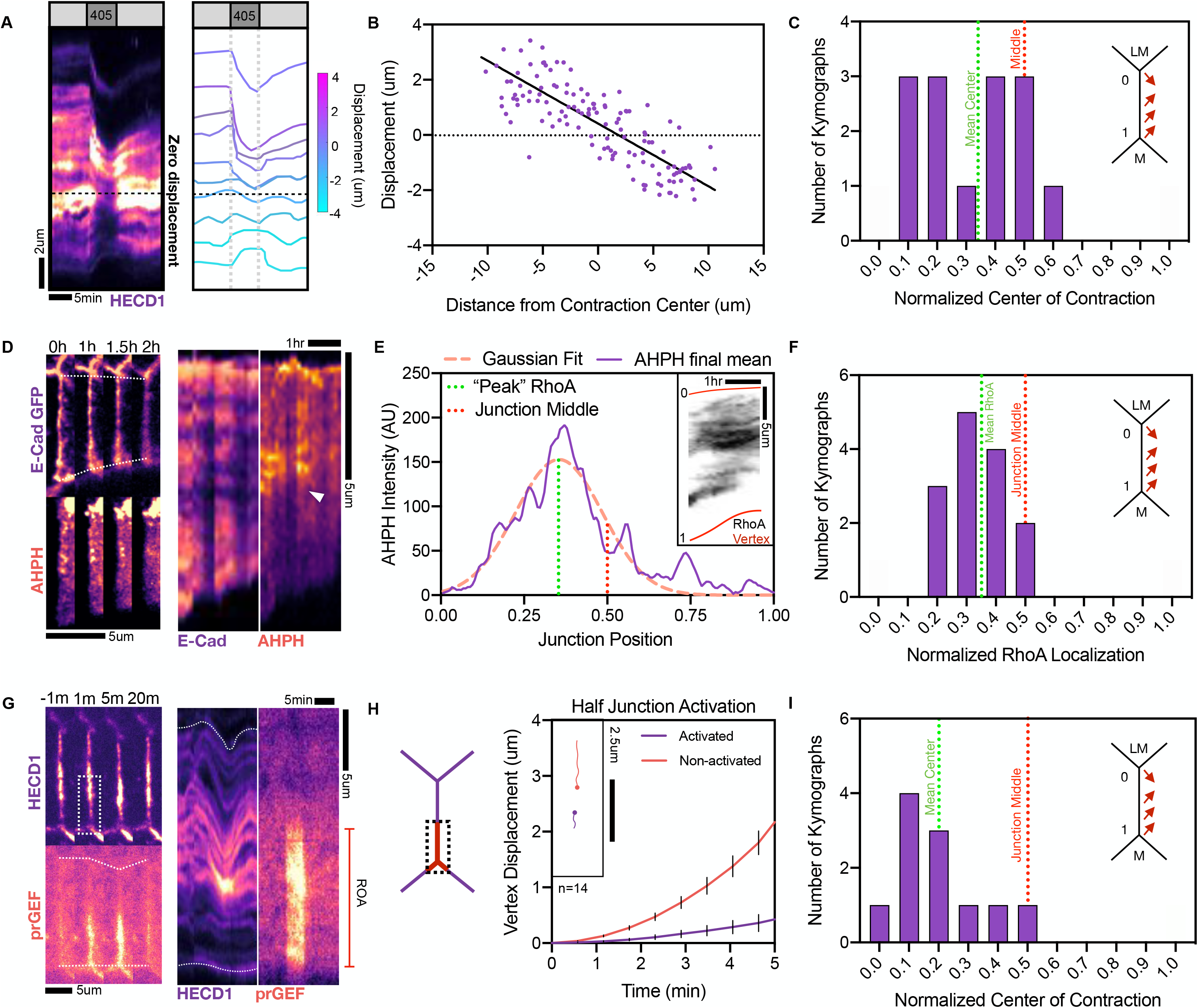
Asymmetric RhoA drives contractile asymmetry. A. (Left) Representative kymograph of an optogenetically activated junction labeled with HECD1 showing asymmetry junction contraction and relaxation. (Right) Fiducial marks seen in the kymograph to the left are color coded according to the amount of displacement within the optogenetic activation period. The location of zero displacement of the fiducial marks is marked with a dashed line. B. Analysis of the displacement of each fiducial mark’s flows as a function of the distance from the contraction center showing linear displacement from one end of the junction to the other, indicating a uniform contraction of the junction. D. Inlay shows diagram of the Less-motile (LM) vertex being labeled as 0 and the Motile (M) vertex being labeled as 1. Red arrows represent the extent of the vertex motion along the junction during contraction. Analysis of the localization of zero displacement (as seen in A) of the fiduciary marks indicates the center of the junction as being skewed towards the less-motile vertex. D. (Left) Representative image of a junction in CN03 treatment expressing E-cadherin-GFP and the transfected RhoA biosensor, AHPH. Junction shows asymmetric contraction with a RhoA flare along the junction. (Right) Kymographs of the image stills shows asymmetric junction contraction and a RhoA flare that is biased towards the less-motile vertex. E. Analysis of the junctional AHPH intensity plots averaged over the last 5 frames of the kymograph (inlay) fitted to a Gaussian curve. Dotted line indicates the peak of the Gaussian fit, indicating the centralized location of the RhoA biosensor. F. Pooled analysis of the peak of the RhoA biosensor, as calculated in E, showing mean junctional RhoA localization as being skewed towards the less-motile vertex. G. Representative image and kymograph of a junction undergoing half-junction activation at the bottom junctional region. H. Vertex displacement analysis of bottom-junction activation showing contractile asymmetry between two vertices. Inlay shows individual vertex tracks for two vertices of the same junction. I. Normalized center of contraction analysis for bottom-junction activation showing the center of contraction is skewed towards the region of activation.

Using these kymographs, we then measured fiducial marks’ and vertices’ maximal displacement over time (Figure 2A). The maximal displacement of each contracting E-cadherin puncta was measured as a function of the position along the junction, for which we then calculated a linear fit. The kymograph’s center of contraction was determined by the root value of that linear fit, and the center of contraction was then normalized so that the position of the more motile vertex was 1 and the less motile vertex 0. If the contraction occurred heterogeneously, the displacement of a fiducial marker would not be proportional to its distance from the contraction center. Instead, we found that the maximum fiducial displacement as a function of the distance from the center of the junction contraction revealed a monotonic, linear trend indicative of a uniform contraction (Figure 2B).

We next sought to characterize the location of the center of contraction along the junction length by identifying the location of zero displacement along the junction length. To compare junctions of varied lengths, we normalized by the junction length. For consistency, we identified the less mobile vertex position as 0 and the more mobile vertex position as 1. For a symmetric contraction, we expected to see the center of contraction at the midpoint of the junction, or 0.5. Instead, we found the contraction center to be skewed towards the less-motile vertex (Figure 2A, 2C). Analyzing multiple kymographs revealed that the mean center of junction contraction was consistently closer to the less-motile vertex with a mean of 0.32 (Figure 2C).

It is plausible that, even though we are stimulating GEF recruitment uniformly along the junction, that RhoA activity is not uniform along the junctions, giving rise to heterogeneous stress along the junction. To explore this, we used the RhoA biosensor (AHPH) containing the GTP-RhoA binding C-terminal portion of Anillin (Piekny and Glotzer, 2008). We transfected the E-cadherin expressing cells with the AHPH and then visualized RhoA activity during a CN03 wash-in experiment. We observed discrete sub-junctional region of heightened active RhoA, which we termed flares (Figure 2D, 2E, Supplemental Movie 3). These flares were absent from junctions without CN03 treatment (Supplemental Figure 3A-F). We measured the intensity of active RhoA along the junction and found that the peak, the central flare location, was skewed towards the less-motile vertex (Figure 2D, 2E inlay, Supplemental Movie 3). Fitting a Gaussian curve to this data, we labeled the peak of this Gaussian as the location of the “peak” RhoA region (Figure 2E). By analyzing fourteen kymographs, we found that the mean RhoA flare position was skewed towards the less-motile vertex, with an average position of 0.35 (Figure 2F).

The above data indicated that the location of RhoA flares were critical in determining asymmetric contraction, with reduced mobility of the vertex proximal to active RhoA. To test this hypothesis, we exploited the optogenetic approach to systematically activate only a portion of the junction. When the lower half of the junction was activated, the junction contracted to about 85% of its original length, similar to the extent for full junction activation. The vertex proximal to the region of activation (ROA) was significantly less mobile than the distal vertex (Figure 2G, 2H, Supplemental Movie 4). Kymograph analysis in the HECD1 channel revealed that the center of contraction for the half junction activation was at the relative position of 0.2 (Figure 2I). Altogether these data indicate that asymmetry in active RhoA dictates the bias in vertex motion.

### Mechanosensitive E-cadherin induces vertex friction at less-motile vertices

RhoA acts at cell-cell interfaces to regulate cell morphology through its effect on actomyosin tension and adhesion strength (Cavanaugh et al., 2020; Levayer et al., 2011). To explore the possibility that changes in adhesion strength underlie vertex immobility, we analyzed E-cadherin localization, as visualized by HECD1 fluorescence, at tricellular vertices during whole junction optogenetic stimulation. We observed HECD1 fluorescence in punctae along the junction and at both vertices. We monitored the HECD1 fluorescence at both vertices during an activation experiment. At the more motile vertex, we found that the HECD1 intensity did not vary significantly during the experiment (Figure 3A, red arrow). By contrast, at the less mobile vertex, we found there was a marked increase of HECD1 immediately after activation which diminished after exogenous stimulation was removed (Figure 3A, 3B, Supplemental Movie 5). This trend was consistent across numerous junctions and paired vertices (Figure 3C). Together these data indicate that changes in tricellular adhesion may contribute to the observed asymmetry.

**Figure 3:**
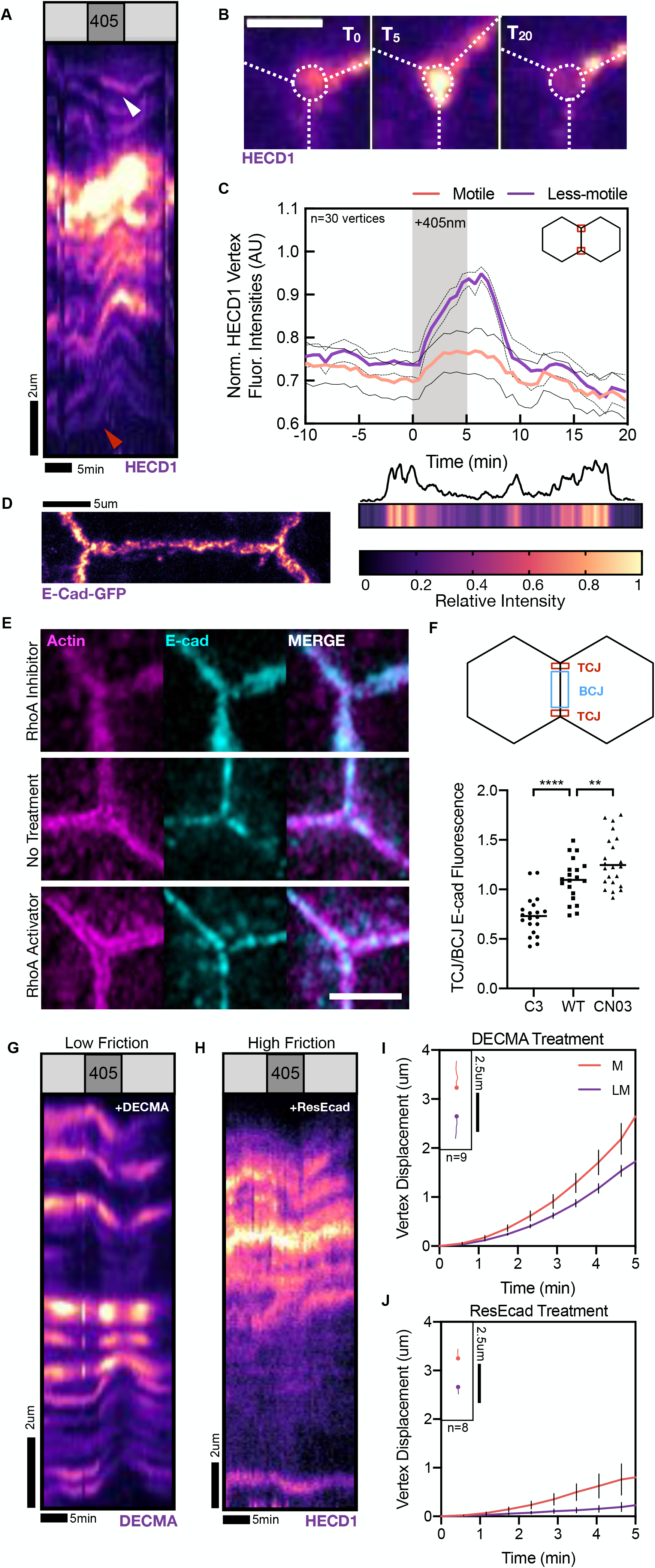
E-Cadherin at tricellular contacts regulates vertex friction. A. Representative kymograph of optogenetic activation showing increases in E-cadherin pooling at the less-motile vertex (white arrow) versus the motile vertex (red arrow). B. Representative image of a less-motile tricellular contact showing E-cadherin pooling at the vertex after 5 minutes of optogenetic activation. Scale bar is 2.5μm. C. Quantification of vertex fluorescence intensities of motile and less-motile vertices. Less-motile vertices show increases in E-cadherin pooling and subsequent vertex fluorescence compared to motile vertices. D. (Left) Representative image of a junction endogenously expressing E-cadherin. (Right) Linescan along the junction for fluorescence intensities and corresponding heatmap showing preferential localization of E-cadherin to the tricellular endpoints. E. (Top) A tricellular junction in C3 transferase treatment stained for Actin (magenta) and E-Cadherin (cyan). (Middle) A tricellular junction in normal media treatment. (Bottom) A tricellular junction with the addition of CN03 RhoA Activator. Scale bar is 2.5um. F. Quantification of the ratio between the tricellular junctions and bicellular junctions’ fluorescence intensities in the different RhoA perturbations. ****=p<0.0001 and **=p<0.05 as calculated by the Student’s t Test. G. Representative kymograph of treated junction with the E-cadherin blocking antibody, DECMA. DECMA treatment introduced symmetry back into contraction upon optogenetic activation. H. Representative kymograph of cells treated with ResEcad showing little-to-no junction contraction. I. Vertex displacement analysis of DECMA-treated junctions showing symmetric contraction. Inlay shows particle tracks of a representative vertex pair during optogenetic activation. J. Vertex displacement analysis of ResEcad-treated junctions showing a severe reduction in the contraction of junctions, resulting in reduced vertex motions and asymmetry. Inlay shows particle tracks of a representative vertex pair during optogenetic activation.

We then visualized the endogenous E-cadherin in our E-cadherin CRISPR cell line to visualize E-cadherin distribution along epithelial junctions. We found preferential localization of E-cadherin at vertices, similar to developmental systems (Figure 3D) (Vanderleest et al., 2018). Upon fixing and staining the tissue for E-cadherin, we saw RhoA-dependent changes in the localization of E-cadherin to tricellular contacts upon RhoA modulation using pharmacological RhoA inhibitors (C3 transferase) and RhoA activators (CN03) compared to wild type (WT) media conditions (Figure 3E, 3F). By inhibiting RhoA activity, we found little tricellular E-cadherin compared to bicellular junction E-cadherin when fixed and stained. In heightened RhoA activity, we found an increase in tricellular E-cadherin recruitment compared to WT controls (Figure 3E, 3F). These data indicate that E-cadherin is mechanosensitive at tricellular vertices, possibly inducing frictional drag at these tricellular vertices.

To explore whether E-cadherin-mediated adhesion acts to impede contraction via its contribution as a source of frictional drag, we next sought to modulate E-cadherin interactions. First, we used a function blocking antibody, DECMA, and its conjugated secondary antibody to visualize junctional dynamics. DECMA binds specifically to EC1 domains on E-cadherin, abolishing any trans interactions between E-cadherin molecules, thereby reducing E-cadherin binding. Upon addition of DECMA, we found a similar labeling pattern of E-cadherin that coated the junction (Figure 3G). Optogenetic activation induced similar junctional contractions compared to WT conditions, but the contraction was more symmetric (Figure 3G, 3I, Supplemental Movie 6) To increase junctional friction, we next sought to increase the levels of E-cadherin through the cell-permeable, pharmacological isoxazolocarboxamide compound, ResEcad (Stoops et al., 2011) (Supplemental Figure 4A). This compound has been shown to induce a dose-dependent increase in E-cadherin levels in adenocarcinoma cells, thereby modulating junctional friction levels. We found ResEcad treatment severely suppressed optogenetically-induced junction contraction (Figure 3H, 3J, Supplemental Fig 4C, Supplemental Movie 7). These data indicate that modulating E-Cadherin levels and interactions, inducing either low or high friction, can influence both the magnitude and asymmetric nature of vertex motions.

### Mechanical model of asymmetric contraction

To quantitatively explain the biomechanical origins of asymmetric vertex motion, we developed a continuum mechanical model for the junction dynamics arising from the balance of tensional forces of the primary junction with the two neighboring shoulder junctions, and a frictional drag acting at the vertices to resist their motion (Figure 4A-B). We modeled the junction as a linear elastic continuum with compressional modulus *E*, tension *Λ*, and dissipating stresses with a friction coefficient *μ*. Our approach stood in contrast to existing vertex models of epithelial tissues (Alt et al., 2017; Farhadifar et al., 2007; Fletcher et al., 2014), where the epithelia are modelled as networks of edges under uniform and constant tension (Noll et al., 2020), with the vertex positions determined by force balance from the neighboring junctions. By modeling the junction as an elastic continuum, we allowed for the junction tension and friction forces to vary along the length of the junction, such that the displacement along the junction would be tracked during a contraction event (Figure 4C). Mechanical force balance at a point along the junction was written as

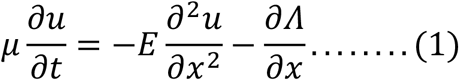

where *u*(*x*, *t*) was the displacement along the junction at time t, and *x* was the position along the junction. The shoulder junctions were modeled as providing a spring-like resistance to motion, with an effective stiffness *k* that depended both on both the tension and the geometry of the shoulder junctions (see Model Methods). Force balance at the tricellular vertices was given by

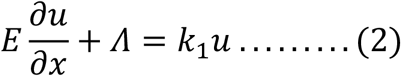

at *x* = *x*_1_ and

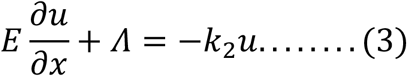

at *x* = *x*_2_, with *k*_1_ and *k*_2_ being the stiffnesses of the two shoulder junctions. To simulate RhoA-induced contraction, we applied a uniform contractile stress for a duration of 5 minutes to a junction initially at rest and recorded the resulting displacements of the two vertices. These displacements were obtained by solving Eq. (1) subject to the boundary conditions given by Eqs (2) and (3). We then used the model to test three different mechanisms for asymmetric vertex motion and heterogeneous mechanical response arising from (i) differential elastic resistance at the shoulder junctions, (ii) differential friction and (iii) asymmetric tension along the junction.

**Figure 4:**
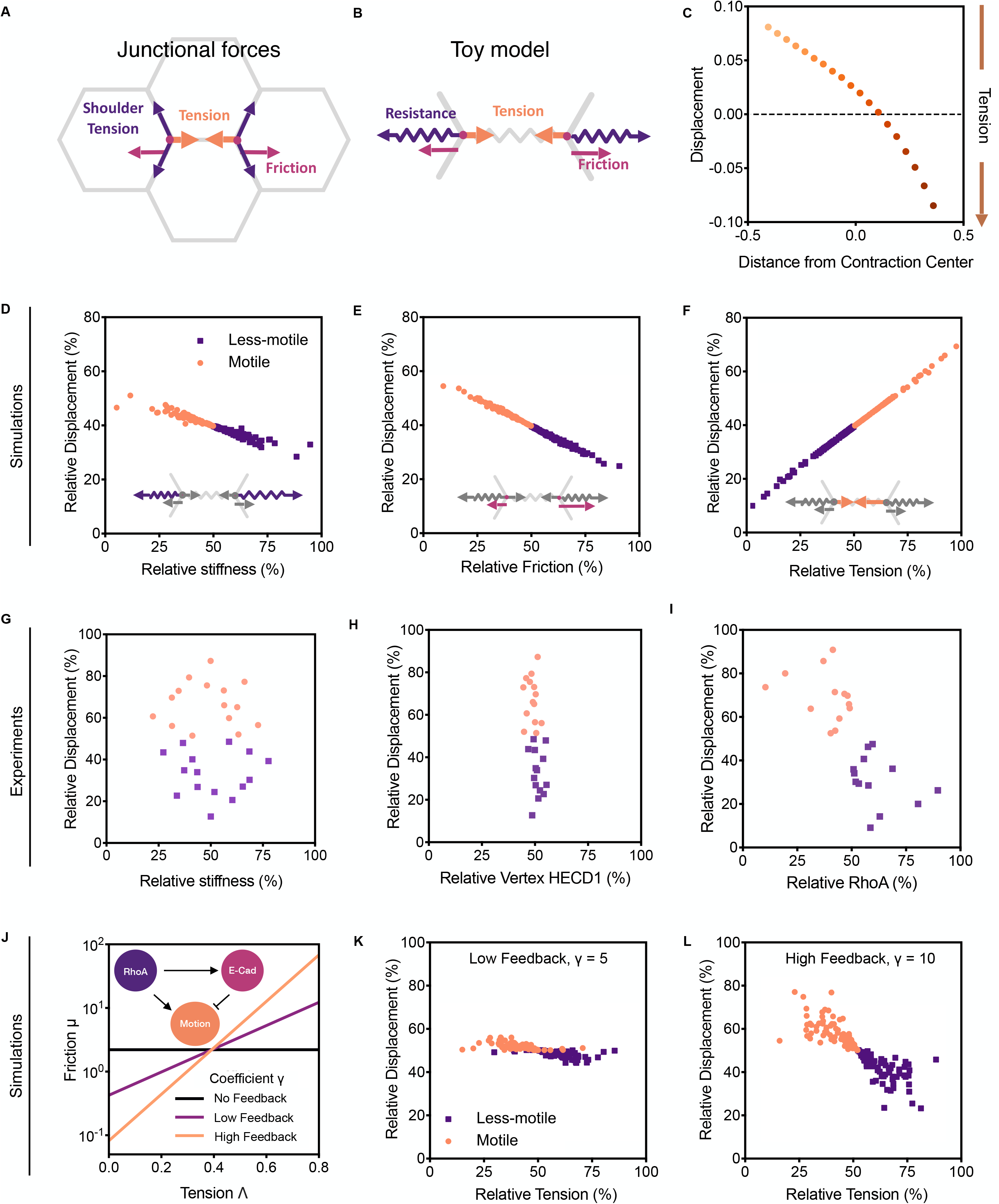
Mechanical forces regulating vertex motion asymmetry. A-B. Toy model schematic, showing the main forces involved in driving vertex asymmetry. Shoulder junctions are modeled through a spring-like resistance to motion of the vertices. The junction is modeled as an elastic continuum, where tension and friction may vary along the junction, having difference values at the two vertices. C. Displacement after contraction against initial position along the junction. D. Simulations of the relative stiffness against relative motion of the whole junction. E. Simulations of the relative friction against the relative motion of the whole junction. F. Simulations of the relative tension at the vertex against the relative motion of the whole junction. G. Experimental data plotting relative motion as a function of relative stiffness showing no correlation. H. Experimental data plotting relative motion as a function of the HECD1 intensity ratio at T-10 before optogenetic activation, providing insight into the relative amounts of friction between the motile vertices and less-motile vertices. I. Experimental data plotting relative motion as a function of the RhoA percentage at each vertex compared to the entire junction. J. Friction against tension for different force scales *γ*. Inlay shows schematic representing feedback between RhoA and E-Cadherin that generates vertex motion. K-L. Percent motion against percent tension for *γ*=5, and *γ*=10.

We first tested how the asymmetry in vertex motion was regulated by differential elastic resistance from the shoulder junctions using our continuum mechanical model. For each vertex, we sampled the shoulder junction stiffness *k_i_* from a Normal distribution with mean *k*_0_ and standard deviation *k*_0_/3. For each vertex, we then compared the percentage of total vertex displacement (relative displacement), *u_i_*/(*u*_1_ + *u*_2_), against the percentage of total shoulder stiffness (relative stiffness), *k_i_*/(*k*_1_ + *k*_2_). Expectedly, we found that vertex displacement depended linearly on shoulder stiffness, with relative displacement decreasing with increasing relative stiffness (Figure 4D).

To test the model predictions using our experimental data, we estimated the elastic resistance at shoulder junctions by computing the tensions along shoulder junctions and change in their geometries during a contraction event, as measured by calculating junction length and the interior angles normal to the activated junction (see Model Methods). From the angles between the activated junction and its neighbors, we calculated the relative tensions on each junction by balancing forces both along the junction and perpendicular to it. From these tensions, we then calculated the differential change in force due to a change in vertex position, which defines the effective stiffness of the shoulder junctions (see Methods). However, when we quantified the relative stiffness using data from our optogenetic experiments, we found no correlation with relative vertex displacement (Figure 4G), indicating that asymmetric elastic resistance at the vertices do not play a role in predicting asymmetric vertex motion upon contraction.

An alternative mechanism for asymmetric vertex motion could arise from heterogeneous adhesive properties at the tricellular vertices or even along the junction proper that may alter the frictional drag. Indeed, our experimental data showed that there is a marked increase in E-cadherin levels at the less motile vertex compared to the motile one during an optogenetic activation (Figure 3A). We therefore sought to test if different frictional forces at the vertices could capture the asymmetric vertex motion. At each vertex, friction was set to a random value sampled from a normal distribution with mean *μ*_0_ and standard deviation *μ*_0_/3, and values were linearly interpolated along the junction. We found a linear dependence of relative displacement on relative friction *μ_i_*/(*μ*_1_ + *μ*_2_), with *μ*_1_ and *μ*_2_ being the friction coefficient at the two vertices, such that increased friction resulted in reduced motion (Figure 4E). As an estimate of the friction in experimental measurements, we measured the relative percentage of HECD1 at each vertex compared to the total amount of HECD1 within each vertex pair. To our surprise, we did not find any correlation between vertex motion and initial cadherin-mediated friction (Figure 4H). Instead, we found that HECD1 intensities were relatively even between each vertex before optogenetic activation.

Finally, we considered the effects of varying tension along the junction induced by RhoA mediated contractility. We varied tension along the junction by setting the tension at each vertex to be a random value sampled from a normal distribution with mean *Λ*_0_ and standard deviation *Λ*_0_/3, and linearly interpolated tension along the junction. We found that vertices under higher tension (more contractility) underwent larger displacements (Figure 4F). To measure relative junction tension, we returned to our CN03 wash in experiments to measure RhoA intensities. We split the junction into two halves and measured the relative intensity of AHPH at each junctional portion compared to the total amount of AHPH along the junction proper. Plotting relative displacement as a function of this percentage of RhoA intensity, we found a correlation between less-motile vertices and the relative amount of junctional RhoA present (Figure 4I). Here, data suggested that the closer the RhoA was to a vertex, the less it moved, consistent with our data in Figure 2G. This was starkly contrasted to highly motile vertices, which were distal to RhoA regions and experienced little RhoA-mediated tension. Together these data suggest that asymmetries in friction, tension, and stiffness parameters alone were insufficient to explain asymmetries in vertex movement during junction contractions.

Our experimental observations informed us that vertices with higher recruitment of RhoA moved less (Figure 2); in contrast, simulations predicted that tension increased proximal vertex displacements (Figure 4F). At the same time, less mobile vertices also showed a marked increase in E-cadherin levels during an optogenetic activation (Figure 3A-C). Thus, there may be force-dependent recruitment of E-cadherin, resulting in increased cell-cell friction. This was conceptually similar to mechanosensitive frictional drag of focal adhesion complexes, where the friction increases with increasing cell-substrate traction stress (Aratyn-Schaus and Gardel, 2010). If this feedback between tension and friction were high enough, then an increase in tension would increase friction to such an extent that these vertices would move slower.

To test this model, we allowed force-dependent friction by tension along the junction in our continuum model. Again, we varied tension along the junction by setting the tension at each vertex to be a random value sampled from the normal distribution with mean *Λ*_0_ and standard deviation *Λ*_0_/3, and linearly interpolated tension along the junction. Using a low-force catch bond model, the friction coefficient was given by *μ* = *μ*_0_*e^Λγ^*, where *μ*_0_ was the friction coefficient at zero tension, and *γ* was a feedback parameter with *γ*^−1^ setting the tension scale for mechanosensitive frictional slip (Figure 4J). For *γ*=0, there was no feedback between tension and friction, such that there is increasing motion with increasing tension (Figure 4F). For intermediate feedback strength, *γ*=5, an increased tension was countered by increased friction, resulting in no correlation between relative junctional tension and displacement (Figure 4K). For high feedback between tension and friction, *γ*=10, the increase in friction was high enough to slow down motion at high tension vertices. As a result, we found a negative correlation between vertex tension and motion (Figure 4L), successfully recapitulating the experimental data (Figure 4I). These simulations, coupled with experimental data, indicated that the E-cadherin recruitment at tricellular vertices likely increased the local friction coefficient to limit junction contraction in a RhoA-dependent manner.

### Local Rho drives E-cadherin pooling

Our data uncovered a positive feedback between RhoA activation and E-cadherin recruitment to tricellular vertices, which increases local friction and reduces vertex mobility. To further determine the functional consequence that RhoA-mediated E-cadherin recruitment had on junction contraction, we used our optogenetic approach to activate the tricellular vertex to witness any feedback at this region. Vertex activation was insufficient to induce junction contraction, with the vertices exhibiting little to no vertex displacement compared to WT full-length activation (Figure 5A, 5B, Supplemental Movie 8). However, tricellular vertex activation did induce a 30% increase in E-cadherin over a 5-minute activation that diminished with removal of the exogenous RhoA activation (Figure 5A, 5C). Moreover, we found that the activated vertex experienced a significant increase in E-cadherin intensities compared to the non-activated vertex (Figure 5C). These data indicated that RhoA activation can increase E-cadherin intensity. We hypothesized this increased concentration arose via RhoA-induced centripetal motion of E-cadherin to cause a local pooling in the region of heightened RhoA.

**Figure 5:**
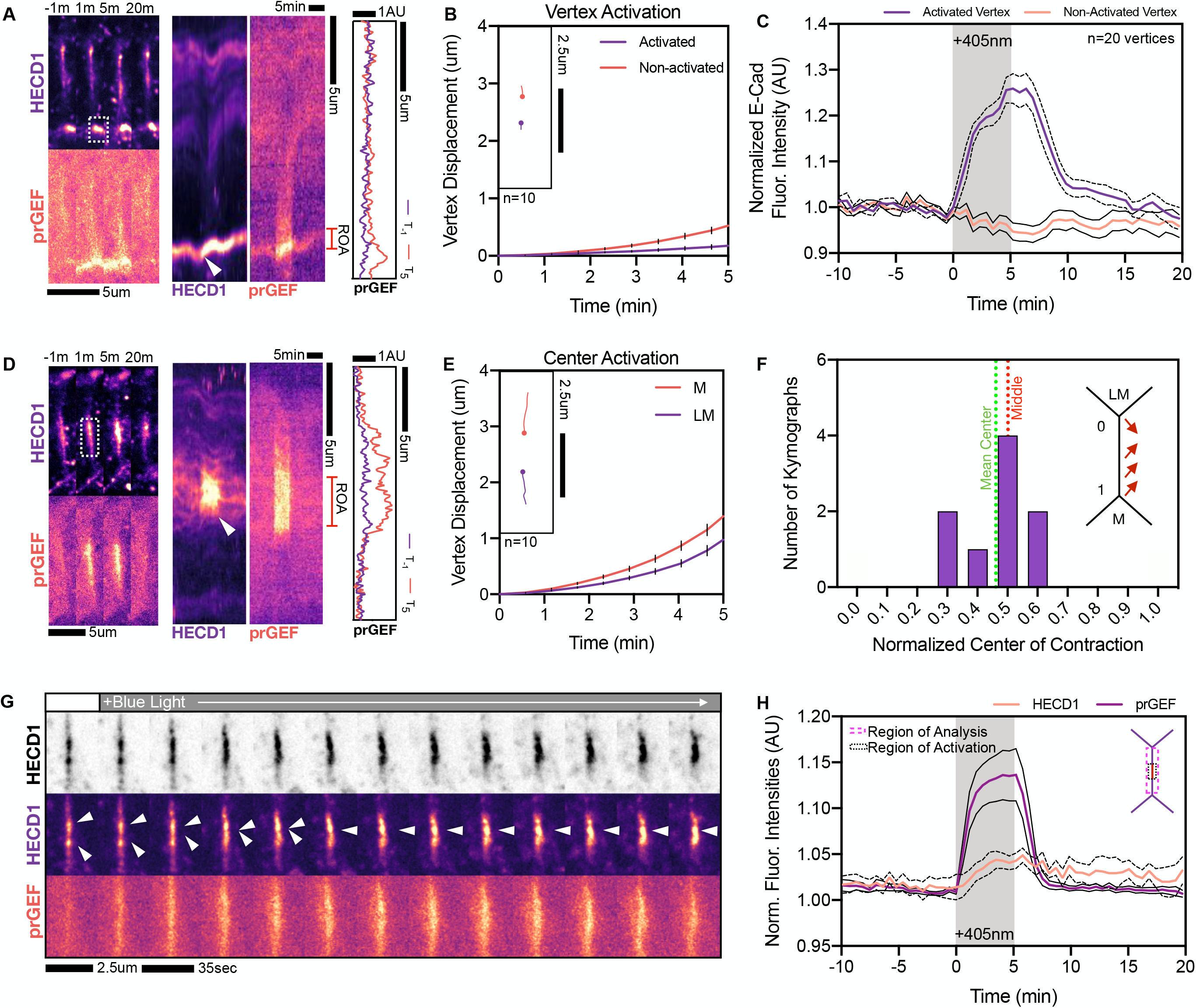
RhoA pools E-cadherin to the location of activation. A. Representative image and kymograph of a junction undergoing only vertex activation at the tricellular contact. B. Vertex displacement analysis of vertex activation showing little-to-no vertex motion within the optogenetic activation period. Inlay shows individual vertex tracks for two vertices of the same junction. C. Normalized HECD1 (E-cadherin) fluorescence intensities for vertices during vertex activation between the activated and non-activated vertices. Activated vertices show increases in E-cadherin fluorescence intensities. D. Representative image and kymograph of a junction undergoing center-junction activation. E. Vertex displacement analysis of center-junction activation showing contractile symmetry is restored. Inlay shows individual vertex tracks for two vertices of the same junction. F. Normalized center of contraction analysis for center-junction activation showing the center of contraction is in the middle of the junction, consistent with where RhoA is activated. G. Representative images of HECD1 and prGEF before and during optogenetic activation, showing a pooling of E-cadherin puncta upon activation (white arrows). H. Quantification of fluorescence intensities of prGEF and HECD1 along the junction seen in G.

To test this hypothesis, we activated only the center of the junction (Figure 5D, Supplemental Movie 8). Activation at the center third of the junction created a contraction whose extent was similar to WT full-length activation. As the center was being activated, there was a noticeable concentration of E-cadherin puncta to the region of activation (Figure 5D). Displacement analysis for the center activation indicated that the contraction was more symmetric, with both vertices moving considerably and relatively evenly upon RhoA activation in the center third of the junction (Figure 5E). However, when the junction was activated in the center third, analysis of the HECD1 fiducial marks revealed that the center of contraction was indeed in the middle of the junction with a mean center of contraction being 0.47 (Figure 5H). These data hint at the possibility that tricellular vertices generate considerable friction during junction contraction, as the lack of activation at vertices produced a symmetric contraction.

To understand the origin of this E-cadherin pooling, we examined junctions activated only at the center third of the junction. We saw E-cadherin pooling upon junctional prGEF recruitment within the activation period (Figure 5G). Here, prGEF recruitment preceded this concentration of E-cadherin, as smaller punctae of E-cadherin coalesced to a concentrated point upon blue light activation (Figure 5G, white arrows, Supplemental Figure 9). We then measured the fluorescence intensities of both the prGEF and the HECD1 along the whole junction without the vertices to exclude the contributions of shoulder HECD1 from adjacent cells. This revealed a 15% increase in prGEF intensities compared to HECD1, whose overall change in intensities along the interfacial junction length was negligible. This led us to a mechanism whereby local, heterogeneous RhoA pulls adhesion molecules from distal regions of the junction to the region of RhoA activation, as analysis of the fluorescence intensities shows a consistent mean fluorescence intensity over time for HECD1 compared to the prGEF (Figure 5H). This suggested a possible mechanism by which E-cadherin slides along contracting actin filaments towards RhoA. No new cadherin was recruited to this region, as fluorescence intensities were conserved along the junction over time.

## Discussion

We present here a new model for vertex mechanoresponse that mediates asymmetric junction contraction via a feedback between RhoA-induced tension and E-cadherin-mediated adhesion. We find that this RhoA-dependent contraction occurs uniformly along the junction length, with both the center of contraction and RhoA localization skewed towards the less-motile vertex. In order to quantitatively model these data, we find that asymmetries in junctional stiffness, friction, and tension parameters alone cannot successfully recapitulate experimental data. Instead, we find experimentally that contraction coincides with the asymmetric pooling of E-cadherin at two paired vertices. The vertex with heightened E-cadherin has increased relative frictional drag, reducing its mobility. We further find that the localization of optogenetic RhoA pools E-cadherin to the region of activation, indicating a novel feedback loop between RhoA and E-cadherin. Incorporating this feedback circuit into our quantitative model, we were able to successfully recapitulate the observed dynamics of vertex asymmetry.

Our study puts forward a novel molecular “clutch” model for tricellular contact engagement during junction contractions. In the absence of RhoA activity, or at distal regions with less RhoA, little E-cadherin is recruited to the vertices. When RhoA-mediated tension is applied to the junction, proximal tricellular adhesions undergo a rapid pooling of E-cadherin that restricts contractile motion in a process similar to that of a “frictional slip” seen in focal adhesions. At focal adhesions, traction stress builds along with frictional drag. The frictional slip is then abrogated once a threshold force is reached, thus providing immobilization of a stable adhesion for adhesion growth (Aratyn-Schaus and Gardel, 2010). We envision a similar mechanism operating at tricellular vertices in that a mechanosensitive rigidity transition of tricellular contacts engages the clutch to strengthen adhesions under load. This adhesion reinforcement restricts vertex motions asymmetrically, as RhoA-mediated tension is stochastically skewed towards one vertex.

These data beg the question as to how RhoA is stochastically placed along the junction. We believe the junction is split into discrete domains that are primed for RhoA activation. These primed regions could be borne out of heterogeneities in adhesive complexes, which exist as puncta along the junction (Cavey et al., 2008). For example, lower junctional E-cadherin levels spatially orient medioapical contractile flows to coordinate junction contractions (Levayer and Lecuit, 2013). We similarly see RhoA flares tracking in regions of low E-cadherin, supporting the notion that E-cadherin-depleted domains could specify primed RhoA regions (data not shown). These domains’ potential for RhoA activation can be exacerbated by the junctional landscape. The local junction composition, specifically lipid and other protein signaling, could generate these distinct contractile units. Indeed, lifetimes of active GTP-RhoA can be enhanced via a coincidence detection scheme upon cyclic binding to the lipid PIP_2_ and the junctional protein Anillin (Budnar et al., 2019). Protein-lipid microdomains, scattered along the junction, could therefore create a permissive environment for RhoA activation that is necessary for junction contractions. Spatial heterogeneities in adhesion, lipids, and protein localization could therefore be critical in determining which portion of the junction is capable of activating RhoA. Further work is needed to discern what specifies these unique microdomains.

These data have serious implications for the canonical mathematical models of epithelial tissues. In traditional vertex models, the tissue is a network of edges and nodes whose geometry and topology depends on active forces. The positions of these vertices anchoring bicellular interfaces are determined by the parameters of interfacial tension and pressure within each cell (Fletcher et al., 2014). Vertices can then move in response to mechanical forces, but the extent of this movement is proportional to the parameters describing vertex friction, shoulder edge tension, and tricellular contact stiffness. Using our heterogeneous junction model, no one single parameter describing friction, tension, or stiffness was able to recapitulate experimental data. Instead, we find that the incorporation of a feedback circuit between RhoA and E-cadherin successfully modeled vertex asymmetry.

Most studies of cell shape changes, to date, concern the movement of bicellular interfaces between two neighboring cells. In development, these junctional zones experience spatially distinct contractile flows that drives qualitatively different and rather opposing junctional responses. Medioapical flows to the bicellular region correspond to junction deformations while flows to the tricellular contacts restrict such contractions (Rauzi et al., 2010; Vanderleest et al., 2018). We see similar junctional responses by optogenetically activating specific junctional zones, with the region of RhoA activation pooling E-cadherin. Our previous work examining stable junction deformations show that longer optogenetic activations facilitate junction length changes through E-cadherin clustering and internalization(Cavanaugh et al., 2020). It would be of interest to see how optogenetic activation of these junctional zones at longer timescales would facilitate their remodeling.

Yet what is the physiological benefit in restricting vertex motion? In the Drosophila Germband, tricellular E-cadherin recruitment is associated with the stabilization of the junctional ratchet. This stabilization ensures progressive interface shortening to facilitate cellular rearrangements(Vanderleest et al., 2018). In our optogenetic system, we do not find stable, irreversible contractions at short timescales but rather reversible junctional deformations. As such, it is unlikely that this vertex reinforcement is sufficient to stabilize junctional shortening. However, it may be necessary to maintain epithelial cohesion under increased tension of neighboring cells. Strong contractions, in principle, could compromise intercellular junctions and barrier functions. Indeed, vertices are principal sites of epithelial fracture in highly tensile epithelia (Acharya et al., 2018). Mechanosensitive reinforcement of vertices could therefore restrict major cell and tissue deformations to maintain tissue homeostasis. This mechanism seems plausible, as RhoA-mediated junctional mechanotransduction is a known regulator of tissue integrity(Acharya et al., 2018).

## Materials and Methods

### Cell culture

E-Cadherin-GFP CRISPR and optogenetic Caco-2 cell lines (generated in Cavanaugh et al., 2020) were cultured in DMEM media (Mediatech, Herndon, VA), and supplemented with 10% Fetal Bovine Serum (Hyclone; ThermoFisher Scientific, Hampton, NH), 2 mM L-glutamine (Invitrogen, Carlsbad, CA), and penicillin-streptomycin (Invitrogen). Cell lines were cultured in a humidified incubator at 37C with 5% CO2.

### Live-cell imaging and transfection

To ensure a confluent and mature epithelial monolayer, Caco-2 cells were plated densely on 2um/ml polymerized collagen gels (unless specified otherwise) coating the bottom of a 4-well Ibidi Chamber (Ibidi). Cells were then allowed to grow for at least 1-2 days to ensure a polarized and confluent monolayer. Ibidi chambers were then placed into a stage incubator with temperature, humidity, and CO2 control (Chamlide TC and FC-5N; Quorum Technologies). All pieces of the stage incubator (stage, adapter, cover, and objective) were maintained at 37C.

To analyze RhoA dynamics, 5ug of AHPH RhoA biosensor DNA (Budnar et al., 2019) was transfected into E-cadherin CRISPR cells using Lipofectamine 3000 (Invitroogen) at least 24 hours before imaging. For CN03 wash-in experiments, cells were imaged in the 488 and 561 channels every 5 or 8 minutes, until 2 hours of timelapse imaging was completed. At the beginning of imaging, either media or 1ug/ml CN03 was added to the media to document junctional responses.

To visualize E-Cadherin in the optogenetic system, we bathed the cells in HECD1 (Abcam) primary and secondary antibody, Alexa Fluor goat Anti-Mouse 647 (Invitrogen), both at a 1:1500 dilution in normal media for at least 24 hours. When applicable, E-cadherin was visualized using DECMA (Abcam) primary and secondary Alexa Fluor Goat Anti-Rat (Invitrogen) antibodies at 1:1500 dilution in normal media for at least 24 hours. Before imaging, cells were washed with PBS and replaced with normal media or media containing chemical perturbations described below. For optogenetic experiments, cells were imaged in the 561 and 647 channel every 35 seconds. The first 10 minutes was to establish a baseline junctional response before the 5-minute optogenetic activation, with the last 15 minutes documenting junctional relaxation. During the activation period, a region around the junction was manually drawn in MetaMorph and adjusted in real time for illumination by the 405nm laser for 1000ms immediately before the acquisition of each image. Laser power was at 1000AU.

For junction and vertex movement analysis, via both CN03 and optogenetic means, we chose to analyze junctions that were distal from cell divisions and/or apoptotic extruding cells to ensure a cohesive monolayer. For picking optogenetic cells, cells were chosen based off of their expression level, which showed junctional recruitment and depletion of the prGEF from the cytosol. All junctions were imaged at the apical plane just below the surface to visualize all vertices and junctional connections.

### Drug Treatments

Cells were treated with a 1:1500 dilution DECMA antibody treatment 24 hours before experimentation. Optogenetic cells were treated with 500uM ResEcad (Calbiochem) or 100uM NSC23766 (Tocris) 24-48 hours before optogenetic activation. WT Caco-2 cells were treated with 1ug/ml CN03 (Cytoskeleton, Inc) or 1μg/ml C3 Transferase (Cytoskeleton, Inc) for at least 4 hours before fixing and staining to analyze E-cadherin localization. E-cadherin CRISPR cells were imaged upon the exposure to 1ug/ml CN03.

### qPCR

Total RNA was isolated with NucleoSpin kits (Macherey-Nagel, #740955). First-strand synthesis was carried out using the SuperScript III system (Invitrogen, 18080-051) with an oligo dT primer and 200 ng of total RNA as input. First -strand reactions were diluted 5-fold and 2 μl was used as template in 20 μl reactions prepared with PrimeTime master mix (IDT,1055772) and PrimeTime pre-designed qPCR primer/probe mixtures from IDT (CDH1: Hs.PT.58.3324071; GAPDH: Hs.PT.39a.22214836). A StepOnePlus instrument (Applied Biosystems) was used for running the qPCR reactions. Relative mRNA levels were determined by the 2-ΔΔCt method utilizing GAPDH as a reference gene.

### Immunofluorescence

Cells were plated onto polymerized collagen gels coating a Lab Tek II Chamber slide (Thermo Fisher Scientific). Once a confluent monolayer was formed, cells were fixed with 4%PFA with 0.1% Triton X-100 in PBS solution (Corning). Permeabilization was achieved through 0.5% Triton X-100 for 10 min and then cells were blocked with 2.5% BSA and 0.1% Triton X-100 in PBS for one hour. Primary antibody, Paxillin at 1:300 or HECD1 at 1:300, was incubated in blocking solution overnight at 4C and then washed at least 3 times for 20 minutes in 0.1% Triton X-100. Slides were the coated with secondary antibody, Alexa Fluor Goat anti-Mouse (Invitrogen), and phalloidin (ThermoFisher) in blocking solution for one hour. After 3 consecutive 20-minute washes in 0.1% Triton X-100, slide chambers were removed and coated with 20ul ProLong Gold (ThermoFisher Scientific). Slides were then sealed with glass coverslips before drying and sealing with nail polish. Slides were then stored at 4C.

### Microscopy

Optogenetic experiments were performed on an inverted Nikon T-E (Nikon, Melville, NY) with a laser merge module with 491, 561, and 642nm laser lines (Spectral Applied Research, Ontario, Canada) with a Yokogawa CSU-X confocal scanning head (Yokogawa Electric, Tokyo, Japan). The Zyla 4.2 sCMOS Camera (Andor, Belfast, UK) collected the images. Optogenetic activation was achieved using a Mosaic digital micromirror device (Andor) attached to a 405nm laser. Images were collected on a 60X 1.2 Plan Apo water (Nikon) objective. MetaMorph Automation and Image Analysis Software (Molecular Devices, Sunnyvale, CA) controlled all hardware. Fix-and-stain and live-cell imaging of CN03 wash-ins were performed on an LSM 980 system with an Airyscan 2 (Zeiss) detector in super resolution-mode with a 63x NA1.4 oil objective (Zeiss). Microscopy software used was the Zen digital imaging suite (Zeiss).

### Image analysis

Vertex displacement and individual vertex traces were acquired by manually tracking each vertex in a vertex pair using the Manual Tracking tool in Fiji (Schneider et al., 2012). Junction lengths were analyzed by manually measuring in each frame the junction length using the free hand line tool in FIJI software. Junction kymographs were generated with a python script written in FIJI to reconstruct user-drawn line segments along the junction proper. Kymographs were made from unregistered image stacks to preserve asymmetry in junction contraction. Linescans of activated regions and E-cadherin along the junction were taken using the Plot Profile tool of a hand-drawn line along the junction in FIJI. Linescans were taken before optogenetic activation and after 5 minutes of activation. Junction intensity profiles were then normalized for the junction length from 0 to 1. Vertex fluorescence HECD1 intensities were calculated by drawing a circle around the vertex region in each frame and measuring the intensities over the time course using the FIJI intensity analysis tool. Contracted length was calculated by dividing the length of the junction at T=5 divided by the length at T=0 during optogenetic activation. Tricellular enrichment of E-cadherin was calculated by taking the ratio of the tricellular E-cadherin region proximal to the vertex contact and dividing it by the bicellular junction length in between two vertices. This intensity measurement was done by drawing a region proximal to the vertex to analyze tricellular E-cadherin using the Intensity analysis tool, and then calculating the intensity by drawing a box region around the bicellular interface. To analyze focal adhesion size and number, the paxillin channel was thresholded and made into a binary mask to calculate the area of focal adhesions within a cell, as indicated by boundary edges seen from apical actin staining. The binary mask was then overlaid onto the paxillin channel to segment the image and calculate the area of paxillin with a threshold of 0.25um^2^ and also the number of focal adhesions within that cellular region identified by apical actin staining. Percent movement was calculated as the displacement of each vertex from the original vertex position in a kymograph divided by the sum of both vertex displacements. Contractile uniformity within each junction was analyzed by manually tracking E-cadherin puncta in each kymograph using the paintbrush tool in FIJI. The maximal displacement of each contracting E-cad puncta as a function of the position along the junction was found and linearly fit using the MATLAB fit function. The kymograph’s center of contraction was determined by the root value of the linear fit, and the center of contraction was then normalized so that the position of the less-motile vertex was 0 and the more motile vertex as 1, meaning the center of the junction would be the position of 0.5. RhoA localization along the junction was found by averaging the AHPH RhoA intensity at the final five timepoints within the kymograph and fitting it to a gaussian using the Matlab fit function. The junction position of the gaussian peak was determined to be the center of RhoA localization and normalized.

### Model methods

The junction is model by an elastic continuum with Young’s modulus E, RhoA induced contractility *Λ*(*x*) and friction *μ*(*x*), which my both vary along the junction. The shoulder junctions are modelled as provided a simple spring like resistance to deformation, with stiffness *k*.

To numerically solve the model for the junction, we discretize the system into *n* equally spaced points along the junction, *x_i_*, with tension *λ_i_* and friction coefficient *μ_i_*. The discretized equations of motion are given by:

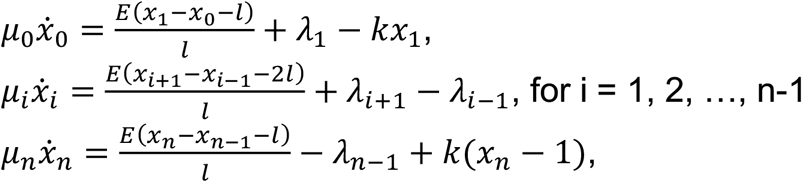

where 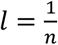 is the distance between position along the junction. The equations are then integrated numerically over time using the python package scipy. For each set of simulations, 100 samples are taken.

The default model parameters are given in the table below. These values are used unless otherwise stated. Parameters are fit by comparing simulations to 5-minute contraction data, and 20-minute contraction data at half-light intensity from Cavanaugh et al, by applying half the tension.

| Parameter | Symbol | Value |
| --- | --- | --- |
| Shoulder stiffness | $k$ | 1 |
| Junction Young's Modulus | $E$ | 1 |
| Friction Coefficient | $\mu$ | 2.5 / min |
| Tension | $\Lambda$ | 0.4 |
| Number of points | $n$ | 21 |

### Shoulder Stiffness Calculations

To estimate the mechanical resistance to motion from the shoulder junctions, we use a simple line tension model of the junctions. Assuming that line tension from the junctions is under force balance, we may calculate the relative tensions from the force balance and derive an effective mechanical energy of the system as the central junction changes length. From this, the second derivative gives us the mechanical stiffness from the shoulder junctions. Let *λ* be the tension of the central junction, *λ*_1_ and *λ*_2_ the tensions of the two shoulder junctions, and *θ*_1_ and *θ*_2_ be the angles between the shoulder junctions and the central junction, and *l*_1_ and *l*_2_ be the initial shoulder junction lengths.

By force balance we have:

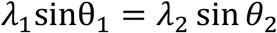

and

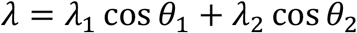

in the x and y directions, respectively, which give the relative tensions.

Next, we calculate the effective resistance from the shoulders by considering the second derivative of the energy with respect to the junction length, *y*. We can write the shoulder junction lengths as

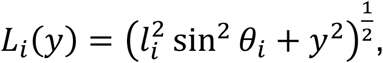

with first derivative

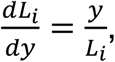

and second derivative

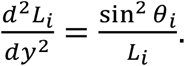

Thus, the mechanical energy *E* = *λ*_1_*L*_1_ + *λ*_2_*L*_2_ − *λy* has second derivative at the initial position of 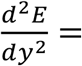 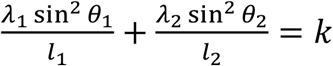, the effective shoulder stiffness.

### Quantification and statistical analysis

Prism software (Graphpad, La Jolla California) was used to establish statistical significance under specific experimental conditions using the two-tailed Student t-test. n represents the number of junctions analyzed in each experiment, which is indicated in the figurers.

## Acknowledgements

KEC acknowledges an HHMI Gilliam Fellowship, National Academies of Sciences Ford Foundation Fellowship, and NIH training grant GM007183. MLG acknowledges funding from NIH RO1 GM104032 and ARO MURI W911NF1410403. This work was partially supported by the UChicago MRSEC, which is funded by the National Science Foundation under award number DMR-1420709. MFS is supported by an EPSRC funded PhD studentship. SB acknowledges funding from the Royal Society (URF/R1/180187) and HFSP (RGY0073/2018). SB and ASY were supported by grants and fellowships from NHMRC Australia (GNT1123816, 1164462, Fellowship1136592) and the Australian Research Council (DP19010287).

---

**Supplemental Figure 1:**
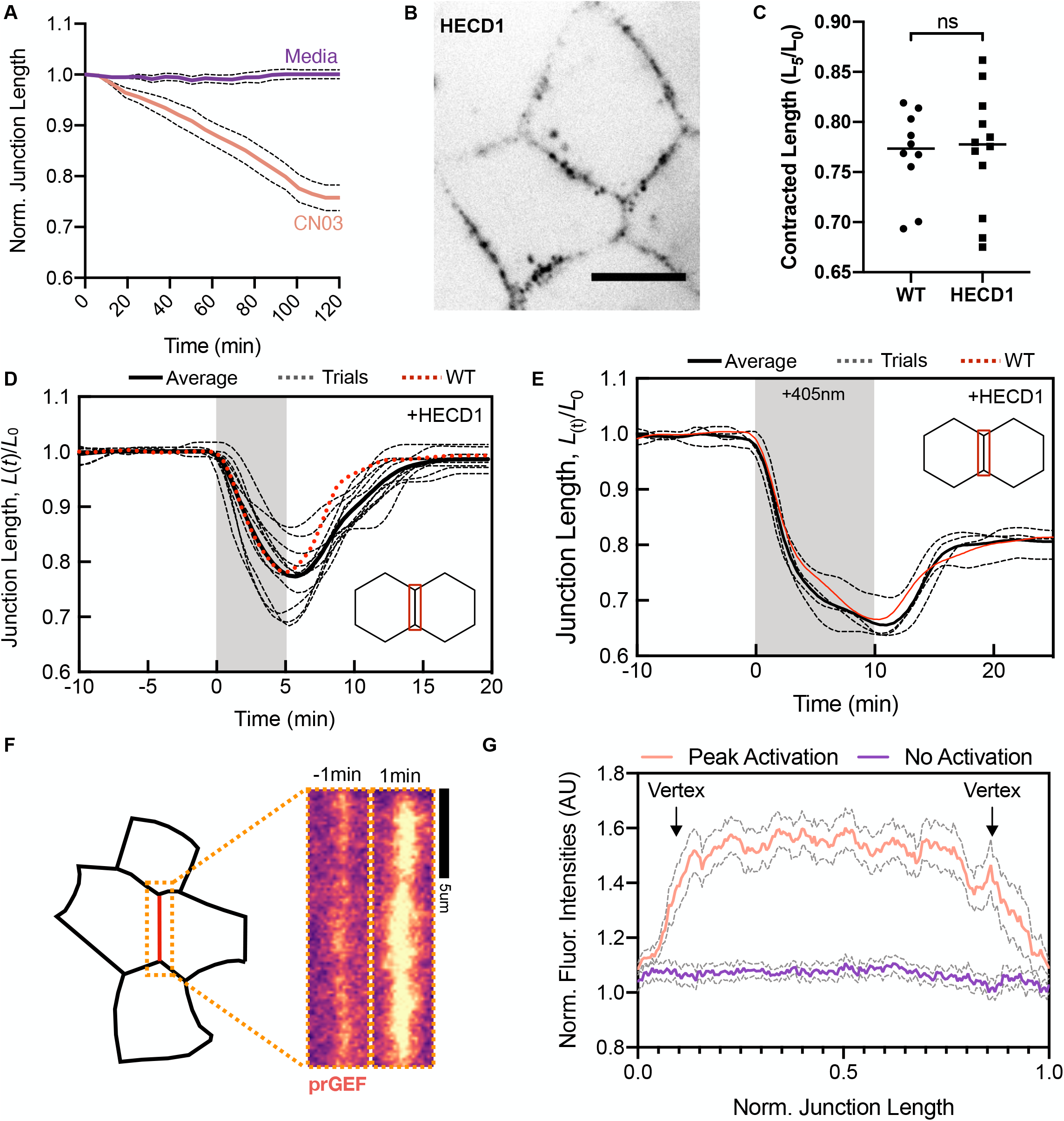
A. Normalized junction length over the course of two hours in WT and CN03 treatment. WT junctions show no junction length changes while CN03 treatment shows junction contraction to about 80% of original lengths. B. Zoomed out image of junctions labeled with HECD1 shows E-cadherin puncta along the junctions. C. Contracted length, or the length after 5 minutes divided by the initial length, after optogenetic activation at T5 between Control and HECD1-labeled cells. HECD1 treatment shows no difference in contracted lengths. D. Normalized junction length over time for each individual junction in HECD1 treatment showing similar contractile phenotypes after 5-minutes of optogenetic activation compared to the average WT junction contraction, with the junction returning to its original length post-activation. E. Normalized junction length for a 10-minute activation of HECD1 treated cells compared to WT controls. Junctions in HECD1 treatment show similar contractile phenotypes as WT conditions, with the junctions readjusting to about 80% of their original lengths post-activation. F. Schematic of an activated junction (left) with uniform junctional prGEF localization before and after optogenetic activation (right). G. Normalized fluorescence intensities along a normalized junction length showing that the peak activation’s prGEF recruitment is uniform along the junction proper compared to pre-activation.

**Supplemental Figure 2:**
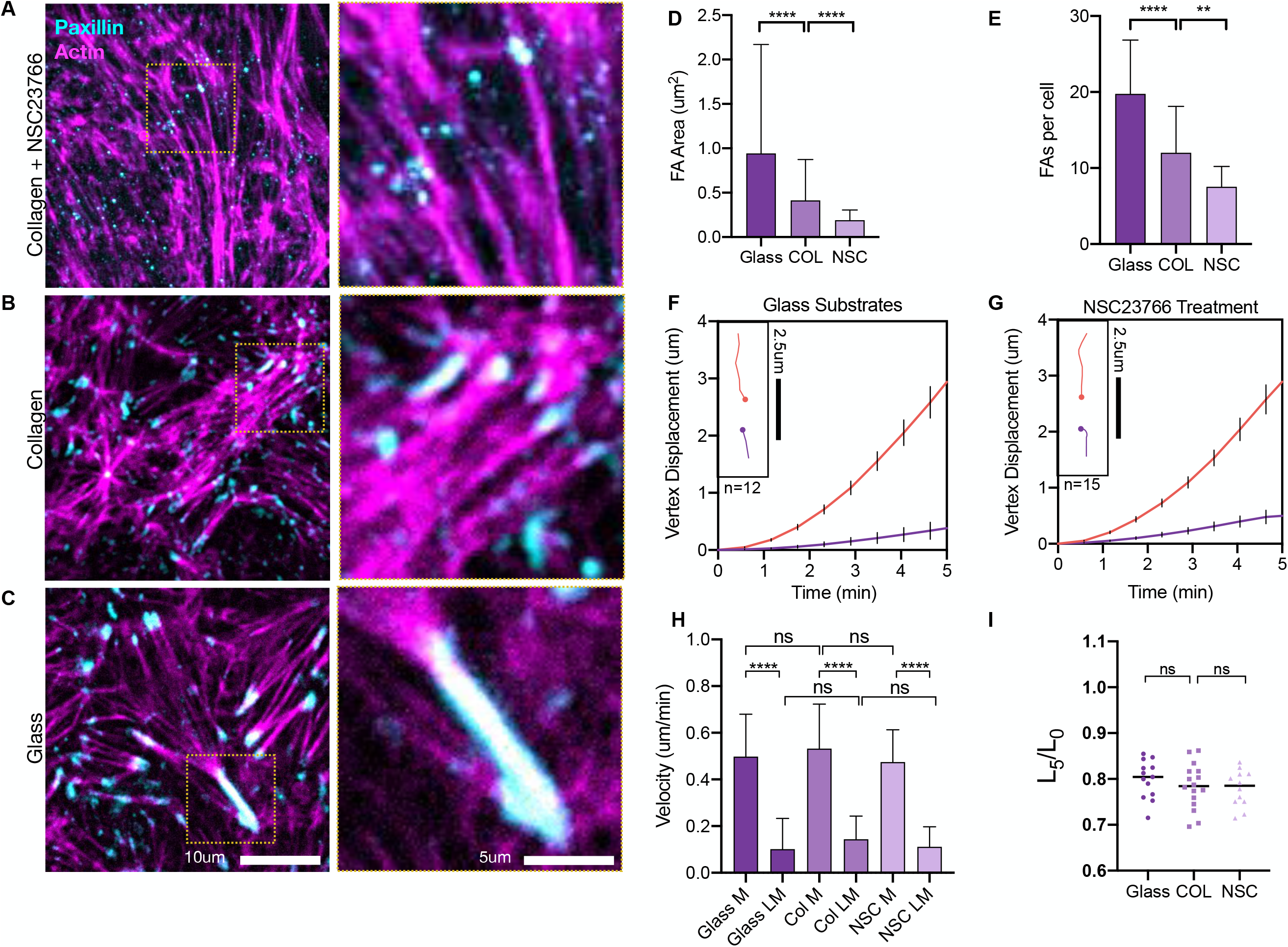
A. Representative image (left) and zoom (right) of the basal substrate surface of cells plated on collagen with Rac Inhibitor, NSC23766, which has been shown to decouple cell-substrate interactions while maintaining cell-cell interactions. Paxillin, a focal adhesion marker, is in cyan and actin is in Magenta. B. Representative image (left) and zoom (right) of the basal substrate surface of cells plated on collagen. Paxillin is in cyan and actin is in magenta. C. Representative image (left) and zoom (right) of the basal substrate surface of cells plated on glass. Paxillin is in cyan and actin is in magenta. D. Quantification of focal adhesion (FA) area among all three conditions showing decreasing FA size with softer substrates (collagen) and substrate decoupling (collagen + NSC23766). Boxes indicate S.D.; whiskers are Standard Error; ****=p<0.0001 as calculated by the Student’s t Test. E. Quantification of FA number per cell among different conditions show decreasing FA number with softer substrates (collagen) and substrate decoupling (collagen + NSC23766). Boxes indicate S.D.; whiskers are Standard Error; ****=p<0.0001; **=p<0.05 as calculated by the Student’s t Test. F. Vertex displacement analysis of activated junctions of cells plated on glass shows vertex asymmetry. Inlays show representative particle tracks of a vertex pair during the optogenetic activation period. G. Vertex displacement analysis of activated junctions of cells plated on collagen + NSC23766 shows vertex asymmetry. Inlays show representative particle tracks of a vertex pair during the optogenetic activation period. H. Analysis of vertex velocities of motile (M) and less-motile (LM) vertices among all three substrate stiffness conditions. Analysis shows asymmetry within each condition, but similar asymmetry across all three conditions tested, suggesting vertex asymmetry is independent of substrate stiffness. Boxes indicate S.D.; whiskers are Standard Error; ****=p<0.0001 as calculated by the Student’s T Test. I. Quantification of the contracted length after optogenetic activation, L5, divided by the initial junction length, L0, among all substrate stiffness conditions tested. Data show the contracted length is similar among all conditions, suggesting substrate does not affect initial junction contractions.

**Supplemental Figure 3:**
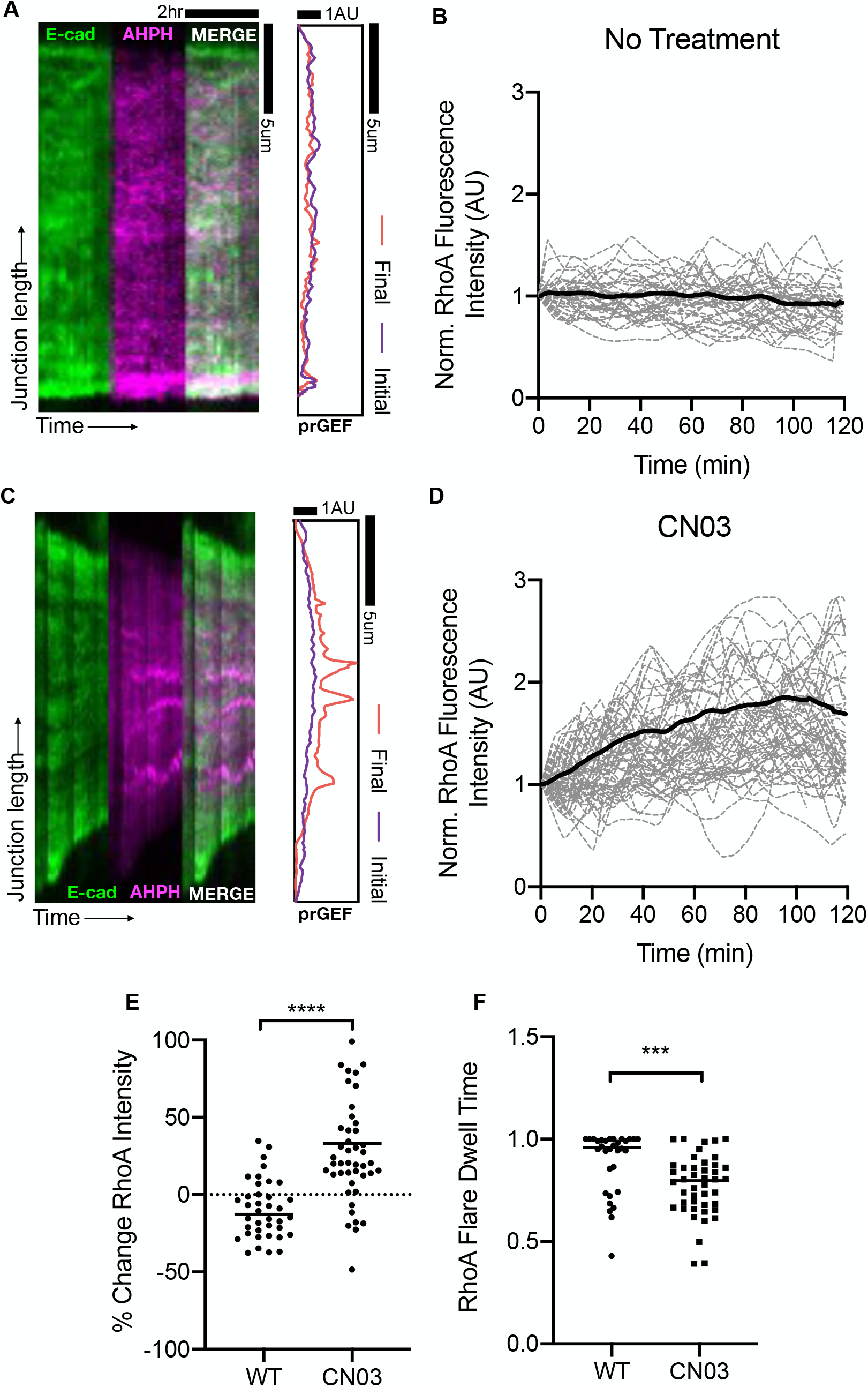
A. Representative kymographs of a junction with no CN03 treatment. B. Normalized RhoA intensity over the course of the 2-hour timelapse. C. Representative kymographs of a junction with CN03 treatment. D. Normalized RhoA intensity over the course of a 2-hour timelapse showing increases in RhoA intensities. E. Percent change in RhoA intensity from the beginning of the kymograph to the end of the kymograph. ****=p<0.0001 as calculated by the Student’s t Test. F. RhoA flare dwell time showing dynamic RhoA flares over the course of the 2-hour timelapse. ***=p0.0009 as calculated by the Student’s t Test.

**Supplemental Figure 4:**
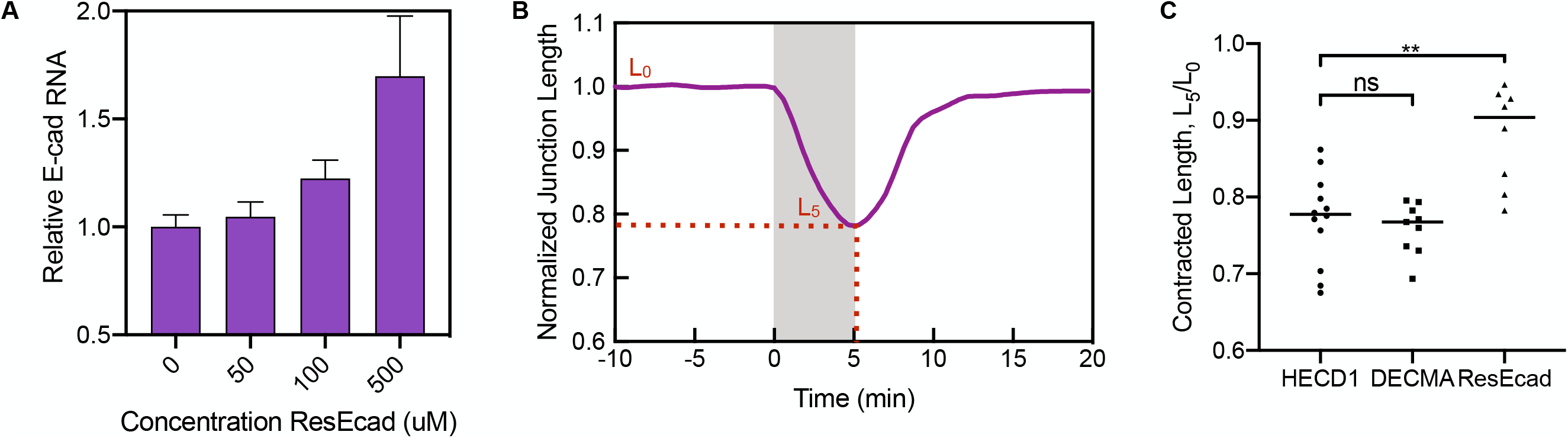
A. qPCR results of relative E-cadherin RNA for 3 replicates as a function of increasing ResEcad treatment. Boxes indicate S.D.; whiskers indicate Standard Error. B. Schematic of normalized junction length with the length at T=0 as L0 and length T=5 is L5. C. Quantification of L5/L0 among different conditions modulating E-cadherin interactions specifically with HECD1, DECMA, and ResEcad. **=p<0.05 as calculated by the Student’s t Test.

## MOVIE LEGENDS

Supplemental Movie 1: Related to Figure 1B, 1C. Fluorescence spinning-disc confocal microscopy timelapse of a (Left) Caco-2 CRISPR E-Cadherin expressing cell junction treated with normal media conditions shows no junction length changes over the course of a 2-hour period. (Right) Caco-2 E-Cadherin expressing cell junction with the 4hr treatment of 1ug/ml CN03 showing asymmetric junction length changes over the course of a 2-hour period. Images taken every 9 min. Scale bar is 5um.

Supplemental Movie 2: Related to Figure 1H, 1I. Fluorescence spinning-disc confocal microscopy timelapse of two Caco-2 TULIP cells expressing the (Top) prGEF and (Bottom) HECD1 labeling showing asymmetric junction length changes within the 5min activation period, as indicated by a white box at T=0. Images taken every 35 seconds. Scale bar is 10um.

Supplemental Movie 3: Related to Figure 2D. Fluorescence spinning-disc confocal microscopy timelapse of a single Caco-2 CRISPR E-cadherin expressing cell junction transfected with the AHPH RhoA biosensor showing asymmetry in contraction and a flare of the AHPH proximal to the less-motile vertex upon 4hr treatment with 1ug/ml CN03. Images taken every 9 min. Scale bar is 10um.

Supplemental Movie 4: Related to Figure 2G. Fluorescence spinning-disc confocal microscopy timelapse of two Caco-2 TULIP cells expressing the (Top) prGEF and (Bottom) HECD1 labeling showing asymmetric junction length changes when 5min activation is only targeted to the bottom region, as indicated by the white box at T=0. Images taken every 35 seconds. Scale bar is 10um.

Supplemental Movie 5: Related to Figure 3B. Fluorescence spinning-disc confocal microscopy timelapse of a Caco-2 TULIP tricellular, low-motile vertex labeled with HECD1 showing HECD1 recruitment to the tricellular contact within the 5min activation region, as indicated by the “+LIGHT.” Images taken every 35 seconds. Scale bar is 2.5um.

Supplemental Movie 6: Related to Figure 3G. Fluorescence spinning-disc confocal microscopy timelapse of two Caco-2 TULIP cells expressing the (Top) prGEF and (Bottom) DECMA labeling showing symmetric junction length changes within the activation period after >24hr treatment with 1:1500 DECMA blocking antibody, as indicated by a white box at T=0. Images taken every 35 seconds. Scale bar is 10um.

Supplemental Movie 7: Related to Figure 3H. Fluorescence spinning-disc confocal microscopy timelapse of two Caco-2 TULIP cells expressing the (Top) prGEF and (Bottom) HECD1 labeling showing a lack of junction length changes within the activation period after 24hr treatment with 500uM ResEcad, as indicated by a white box at T=0. Images taken every 35 seconds. Scale bar is 10um.

Supplemental Movie 8: Related to Figure 5A, 5D. Fluorescence spinning-disc confocal microscopy timelapse of a (Left) Caco-2 TULIP junction expressing HECD1 on the left and the prGEF on the right. The tricellular vertex is activated, as indicated by the white box at T=0. (Right) A Caco-2 TULIP junction expressing HECD1 on the left and the prGEF on the right. The center third of the junction is activated for 5min, as indicated by the white box at T=0. Images taken every 35 seconds. Scale bar is 10um.

Supplemental Movie 9: Related to Figure 5G. A zoomed in fluorescence spinning-disc confocal microscopy timelapse of a Caco-2 TULIP junction activated for 5min at the center third of the junction expressing (Left) HECD1 and (Right) prGEF. Activation at T=0 precedes the pooling of HECD1 punctae to the region of activation. Images taken every 35 seconds. Scale bar is 2.5um.

